# Modeling antimicrobial cycling and mixing: Differences arising from an individual-based versus a population-based perspective

**DOI:** 10.1101/081067

**Authors:** Hildegard Uecker, Sebastian Bonhoeffer

## Abstract

**Background:** In order to manage bacterial infections in hospitals in the face of antibiotic resistance, the two treatment protocols “mixing” and “cycling” have received considerable attention both from modelers and clinicians. However, the terms are not used in exactly the same way by both groups.

**Objectives:** We aim to investigate a model that comes closer to clinical practice and compare the predictions to the standard model.

**Methods:** We set up two deterministic models, implemented as a set of differential equations, for the spread of bacterial infections in a hospital. Following the traditional approach, the first model takes a population-based perspective. The second model, in contrast, takes the drug use of individual patients into account.

**Results:** The alternative model can indeed lead to different predictions than the standard model. We provide examples for which in the new model, the opposite strategy maximizes the number of uninfected patients or minimizes the rate of spread of double resistance.

**Conclusions:** While the traditional models provide valuable insight, care is needed in the interpretation of results.

## Introduction

With bacteria evolving resistance, potent drugs turn ineffective against bacterial infections. Evolution of resistance to a single antibiotic is often rapid, calling for treatment strategies that involve the employment of more than one drug. Besides the simultaneous prescription of a combination of antibiotics to a single patient, several antibiotics belonging to different classes can be used across a community in order to manage the spread of resistance at a population level. This is particularly feasible within a hospital, where drug usage can be controlled and coordinated. Two generic strategies – cycling and mixing of antibiotics – have attracted considerable attention for the phase of empirical therapy.^1–3^ With cycling, drug A and drug B get alternated. With mixing, one half of all patients receive drug A and the other half drug B. Both strategies increase heterogeneity in selection. Assessing the usefulness of these control measures by means of clinical trials is crucial but inherently difficult. While the number of empirical studies is increasing, the overall picture remains inconclusive.^4–8^ Mathematical models therefore remain a helpful tool in understanding which strategy is more promising.^1, 2, 9–17^ Following Bonhoeffer et al. (1997),^1^ a tradition has established of how mixing and cycling are modeled mathematically (for exceptions, see Kouyos et al. (2011)^13^ and Abel zur Wiesch et al. (2014)^15^). It turns out that the verbal description of this implementation is easily misinterpreted. In particular, it differs from empirical practice.

The overall modeling approach taken in all studies is a type of epidemic model with transmission between patients, where infected patients are divided up into several compartments according to the infecting bacterial strain. The traditional implementation of treatment protocols neglects the dynamics that arise if the perspective of individual patients is taken into account. For cycling, this means that when the time for a switch of the drug has come, a patient who is already treated gets switched to another antibiotic rather than that the new drug is only applied to newly infecteds. For mixing, it means that in *every* compartment at *any* time, one half of all patients are treated with drug A and the other half with drug B. Importantly, this is *not* the same as prescribing a random drug to every patient upon infection. To see this, imagine a patient who is infected with a bacterial strain resistant to drug A. Drug B successfully suppresses the infection, leading to a fast recovery. In contrast, drug A is ineffective and the patient recovers slowly. As a consequence, the fraction of patients that get treated with drug A among all those that are infected with an A-resistant strain quickly rises to more than 50% even if newly infecteds have a 50-50 chance to get treated with drug B. The common modeling approach probably comes closest to a situation where at every drug intake, patients take a random drug (although this would have effects at the within-host level that are not captured by the models).

In this article, we set up an alternative model where cycling and mixing refer to the individual antibiotic prescription. We demonstrate by means of examples that this can lead to qualitatively different predictions.

## Methods

We use a model for the spread of bacterial infections in a hospital that is based on a mix of the models in Bonhoeffer et al. (1997) and Bergstrom et al. (2004).^1, 2^ Patients can be uninfected or infected with one of four bacterial strains and can either be already infected at hospitalization or acquire the infection from other patients within the hospital. Resistance can be brought into the hospital from the outside or arise de-novo.

We denote the number of uninfected individuals by *X*. Patients that are infected by the sensitive or a resistant strain are denoted by *S* and *R*_•_, respectively, and the subscript indicates to which antibiotic(s) the strain is resistant. Patients enter the hospital at a total rate of *n*_tot_*µ* and leave at a per-capita rate of *µ* independently of infectious status. In other words, the model assumes that the infection does not increase mortality, and that the infection is not the cause for the hospitalization. *m*_•_ is the fraction of incoming patients for the respective compartment. The transmission probability between uninfecteds and infected patients carrying the sensitive strain is given by *β*. The transmission probability is reduced by a factor (1 – *c*_•_) for infection with a resistant strain; this accounts for a cost of resistance. If a patient infected with a strain sensitive to the currently applied drug gets infected by a resistant strain, the resistant strain can replace the sensitive strain. This happens with a lower probability than infection of an uncolonized patient (reduction by a factor *σ*). Independently of treatment, the immune response of patients leads to recovery at rate *γ*. Treatment with a working drug leads to recovery at rate *τ*. Finally, bacteria can evolve resistance. During treatment with drug A/B, resistance evolves at rate *ν*_*A*_/*ν*_*B*_. Additionally, the sensitive strain becomes resistant to both drugs within a single step at rate *ν*_*AB*_.

In the traditional model, a fraction *χ*_*A*_ is treated with drug *A* and a fraction 1 – *χ*_*A*_ with drug B in *every* compartment at *any* time. Overall, we obtain the following set of differential equations:

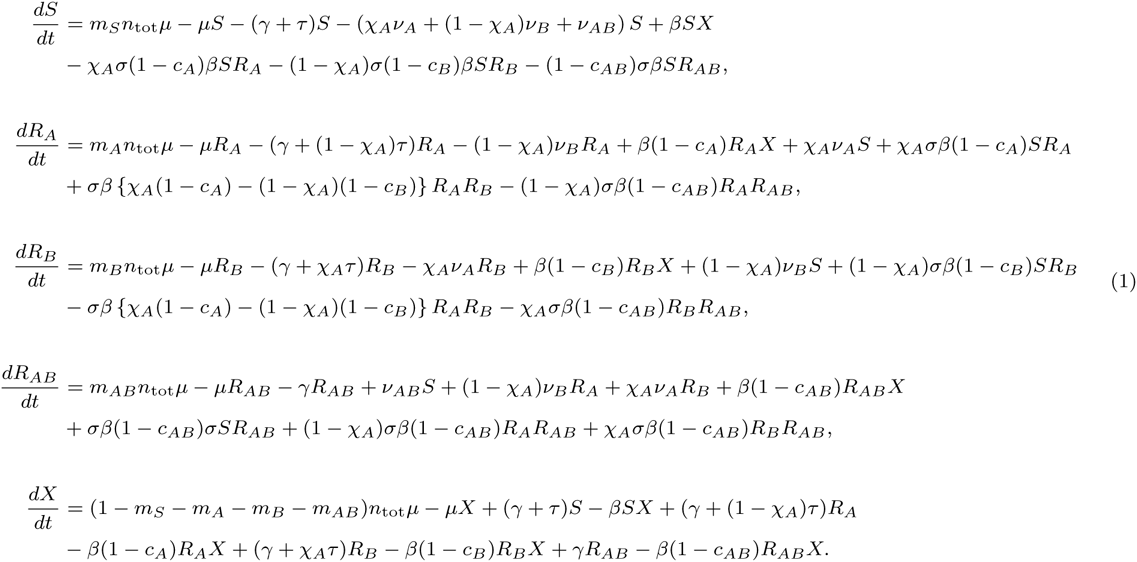

For the alternative model, we split up the compartments *S*, *R*_*A*_, and *R*_*B*_ according to the treatment that patients receive.^13, 15^ The drug is indicated by a superscript. E.g. 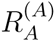 denotes the number of all patients that are infected by a strain resistant to drug A and that are treated with drug A. *χ*_*A*_ is now the fraction of patients who receive drug *A* throughout the entire course of their therapy. This yields:

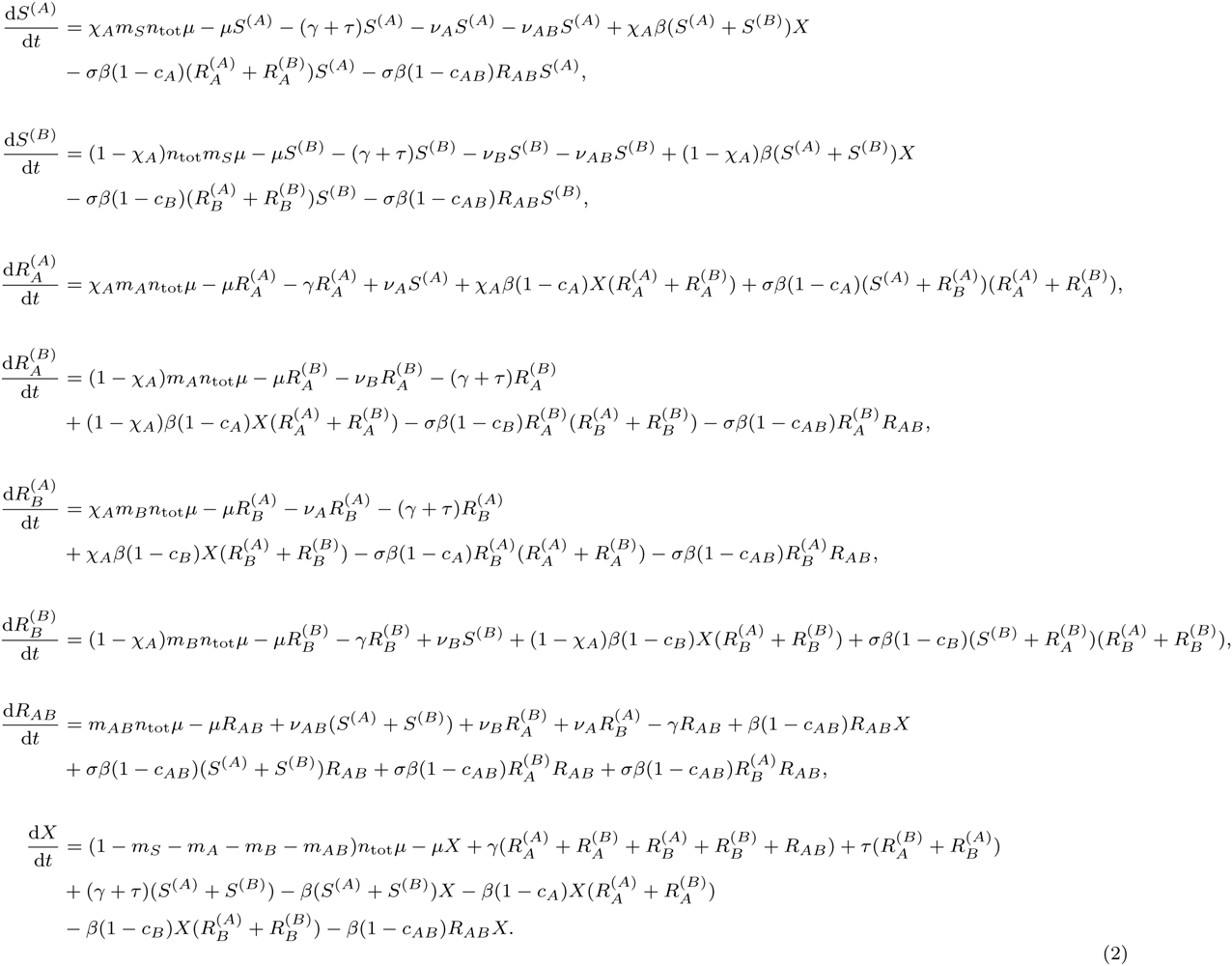

We numerically integrate Eq. (1) and Eq. (2) using Mathematica version 10.4.1.0 (Wolfram Research, Champaign, USA). Integration in R version 3.2.3 using the deSolve package only yields unimportant differences.

## Results and Discussion

We consider two complementary scenarios.

### Absence of double resistance

The de-novo emergence of resistance within the hospital can be neglected (*ν*_*A*_ = *ν*_*B*_ = *ν*_*AB*_ = 0), and no double resistant cases are admitted to the hospital (*m*_*AB*_ = 0). This corresponds to the situation considered in Bergstrom et al. (2004)^2^ and we use the same parameter set; cf. also the similar “Case III” in Bonhoeffer et al. (1997).^1^ We assess the success of the respective strategy by the number of uninfected patients (Figure 1). For both strategies, the number of uninfected patients is lower in the alternative than in the traditional model, since the latter essentially exposes a fraction of patients to both drugs during the course of treatment. Figure 1C shows that in the standard model, cycling performs worse than mixing for all cycling periods,^2^ whilst in the alternative model, we find an optimal cycling frequency around which cycling is slightly better than mixing (cf. also Panels A and B). In this regime, a large fraction of new infections is caused by patients who get still treated with the previous drug and carry the corresponding resistant strain; these new infections are now successfully treated. A similar result as in Figure 1C has been found before in a more complex model, where it was attributed to drug adjustment in case of resistance.^15^

**Figure 1:**
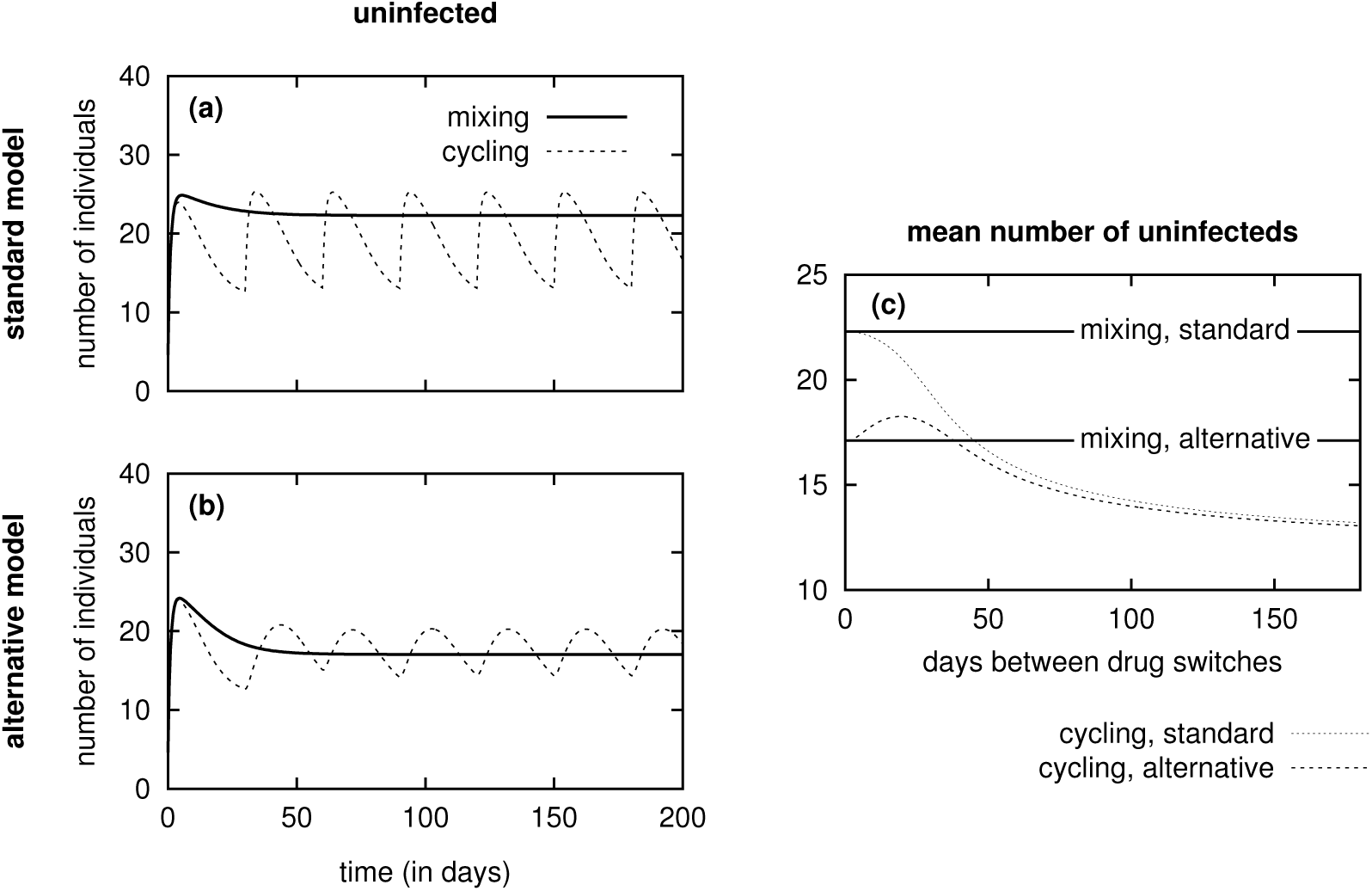
Panels A+B: Number of uninfected patients as a function of time. For cycling, the drug is switched after 30 days. Panel C: Average number of uninfected patients in one cycling period after the transient behavior has decayed. In the standard model, mixing outperforms cycling for all cycling frequencies. In the alternative model, conversely, cycling outcompetes mixing by a small margin for intermediate cycling. Parameters: *n*_tot_ = 100, *m*_0_ = 0.7, *m*_1_ = *m*_2_ = 0.05, *m*_3_ = 0. *µ* = 0.1, *γ* = 0.03, *τ* = 0.25, *ν*_*A*_ = *ν*_*B*_ = *ν*_*AB*_ = 0, *σ* = 0.01, *c*_*A*_ = *c*_*B*_ = *c*_*AB*_ = 0, *β* = 0.01 (see Figure 3 in Bergstrom et al. (2004)^2^). The initial frequencies are given by the equilibrium frequencies in the absence of treatment, *X*_0_ *≈* 4, *S*_0_ *≈* 78, *R*_*A*_ = *R*_*B*_ = 9, *R*_*AB*_ = 0.

### De-novo emergence of resistance within the hospital

There is no influx of patients harbouring resistant bacteria (*m*_*A*_ = *m*_*B*_ = *m*_*AB*_ = 0), and all resistance emerges de-novo within the hospital (cf. “Case II” in Bonhoeffer et al. (1997)^1^). We now consider the spread of double resistance. The picture is essentially the reverse of the medal from the previous paragraph. In the new model, the number of patients infected by a single resistant strain rises to high numbers under the mixing strategy since patients that get treated with an ineffective drug only recover very slowly. At the same time, the colonizing strain is not confronted with the other drug until a new patient gets infected and treated with the second drug. The selection pressure for the double resistant strain is hence low and the spread of double resistance is delayed compared to the standard model (cf. upper and lower Panels in Figure 2). The dynamics under cycling converge to those under mixing for rapid cycling (Figure 2C and F). For slow cycling, the spread of double resistance in the two models is more similar (Panels A and D). For intermediate cycling (Panels B and E), the prediction under which treatment strategy the double resistant strain spreads faster gets reversed.

**Figure 2:**
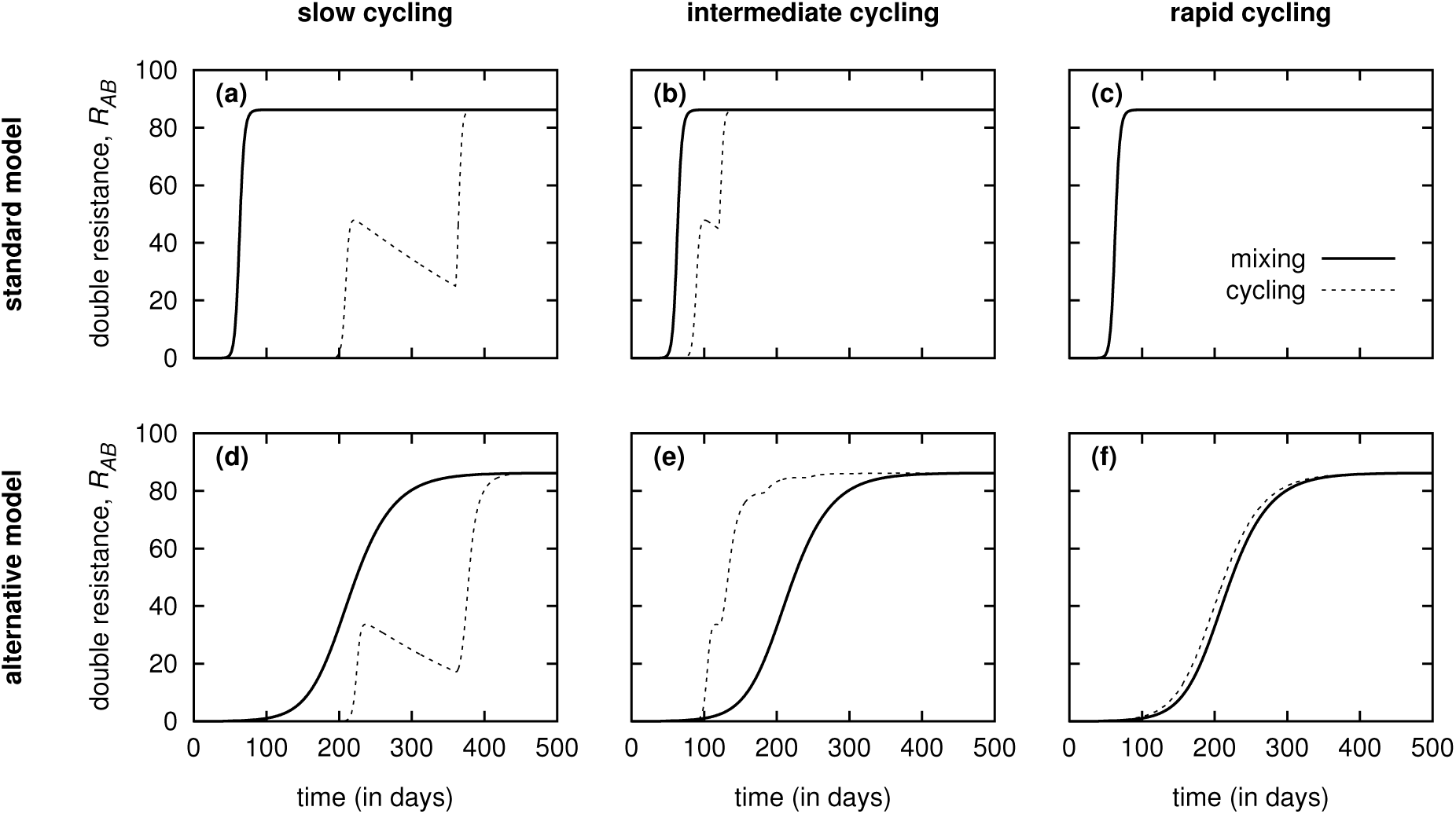
Number of patients infected with the double resistant strain, *R*_*AB*_, as a function of time. For slow cycling (drug switch after 180 days), double resistance spreads more rapidly under mixing than under cycling in both models. For rapid cycling (drug switch after 3 days), the dynamics under the two strategies converge (In Panel C, the lines are indistinguishable). For intermediate cycling (drug switch after 60 days), the predictions of the models differ. While double resistance spreads more rapidly under mixing in the standard model, it spreads more rapidly under cycling in the alternative model. Parameters: *n*_tot_ = 100, *m*_0_ = 0.5, *m*_1_ = *m*_2_ = *m*_3_ = 0, *µ* = 0.05, *γ* = 0.02, *τ* = 0.5, *ν*_*A*_ = *ν*_*B*_ = 10^−5^, *ν*_*AB*_ = 10^−13^, *σ* = 0.01, *c*_*A*_ = *c*_*B*_ = 0.1, *c*_*AB*_ = 0.2, *β* = 0.01. The initial frequencies are given by the equilibrium frequencies in the absence of treatment, *X*_0_ *≈* 4, *S*_0_ *≈* 96, *R*_*A*_ = *R*_*B*_ = *R*_*AB*_ = 0.

### Conclusions

The terms “mixing” and “cycling” do not refer to the same protocol in mathematical standard models and in clinical practice, the former neglecting the behavior of individual patients. In large parts of the parameter space, the results from the standard model and the alternative model that comes closer to the real-life implementation are qualitatively – and sometimes even quantitatively – similar. However, the examples given in this article show that they can also lead to different conclusions, e.g. with the new model, the number of uninfected patients can be lower under cycling than under mixing, where the standard model predicts the opposite. Given such important discrepancies, it seems to be time to be more critical about the established model simplification.

## Acknowledgments

The authors thank Roger Kouyos for helpful discussions.

## Funding

This work was supported by the European Research Council (grant number 268540).

## Transparency declarations

None to declare.

## References

1 S. Bonhoeffer, M. Lipsitch, and B.R. Levin. Evaluating treatment protocols to prevent antibiotic resistance. Proceedings of the National Academy of Sciences, 94:12106–12111, 1997.

2 C.T. Bergstrom, M. Lo, and M. Lipsitch. Ecological theory suggests that antimicrobial cycling will not reduce antimicrobial resistance in hospitals. Proceedings of the National Academy of Sciences, 101(36):13285–13290, 2004.

3 P.J. van Duijn and M.J.M. Bonten. Antibiotic rotation strategies to reduce antimicrobial resistance in Gram-negative bacteria in European intensive care units: study protocol for a cluster-randomized crossover controlled trial. Trials, 15:277, 2014.

4 E.M. Brown and D. Nathwani. Antibiotic cycling or rotation: a systematic review of the evidence of efficacy. Journal of Antimicrobial Chemotherapy, 55:6–9, 2005.

5 J.-A. Martínez et al. Comparison of antimicrobial cycling and mixing strategies in two medical intensive care units. Critical Care Medicine, 34(2):329–336, 2006.

6 A. Sandiumenge et al. Impact of diversity of antibiotic use on the development of antimicrobial resistance. Journal of Antimicrobial Chemotherapy, 57(6):1197–1204, 2006.

7 R. G. Masterton. Antibiotic heterogeneity. International Journal of Antimicrobial Agents, 36(S3):S15–S18, 2010.

8 S. Sarraf-Yazdi, M. Sharpe, K.M. Bennett, T.L. Dotson, D.J. Anderson, and S.N. Vaslef. A 9-year retrospective review of antibiotic cycling in a surgical intensive care unit. Journal of Surgical Research, 176(2):e73–e78, 2012.

9 B.R. Levin and M.J.M. Bonten. Cycling antibiotics may not be good for your health. Proceedings of the National Academy of Sciences, 101(36):13101–13102, 2004.

10 T.C. Reluga. Simple models of antibiotic cycling. Mathematical Medicine and Biology, 22:187–208, 2005.

11 H.-R. Sun, X. Lu, and S. Ruan. Qualitative analysis of models with different treatment protocols to prevent antibiotic resistance. Mathematical Biosciences, 227:56–67, 2010.

12 C.H. Chan, C.J. McCabe, and D.N. Fisman. Core groups, antimicrobial resistance and rebound in gonorrhoea in North America. Sexually Transmitted Infections, 2011.

13 R.D. Kouyos, P. Abel zur Wiesch, and S. Bonhoeffer. Informed switching strongly decreases the prevalence of antibiotic resistance in hospital wards. PLoS Computational Biology, 7(3):e1001094, 2011.

14 U. Obolski and L. Hadany. Implications of stress-induced genetic variation for minimizing multidrug resistance in bacteria. BMC Medicine, 10(89), 2012.

15 P. Abel zur Wiesch, R. Kouyos, S. Abel, W. Viechtbauer, and S. Bonhoeffer. Cycling empirical antibiotic therapy in hospitals: meta-analysis and models. PLoS Pathogens, 10(6):e1004225, 2014.

16 E.M. Campbell and L. Chao. A population model evaluating the consequences of the evolution of double-resistance and tradeoffs on the benefits of two-drug antibiotic treatments. PLoS ONE, 9(1):e86971, 2014.

17 U. Obolski, G.Y. Stein, and L. Hadany. Antibiotic restriction might facilitate the emergence of multi-drug resistance. PLoS Computational Biology, 11(6), 2015.

